# Action-outcome knowledge dissociates from behavior in obsessive-compulsive disorder following contingency degradation

**DOI:** 10.1101/245944

**Authors:** Matilde M. Vaghi, Rudolf N. Cardinal, Annemieke M. Apergis-Schoute, Naomi A. Fineberg, Akeem Sule, Trevor W. Robbins

## Abstract

Goal-directed and habitual systems orchestrate action control. In disorders of compulsivity, their interplay seems disrupted and actions persist despite being inappropriate and without relationship to the overall goal. We manipulated action–outcome contingency to test whether actions are goal-directed or habitual in obsessive-compulsive disorder (OCD), the prototypical disorder of compulsivity, in which prominent theories have suggested that dysfunctional beliefs underlie the necessity for compulsive actions.

OCD patients responded more than controls when an action was causally less related to obtaining an outcome, indicating excessive habitual responding. Patients showed intact explicit action–outcome knowledge but this was not translated normally into behavior; the relationship between causality judgment and responding was blunted. OCD patients’ actions were dissociated from explicit action-outcome knowledge, providing experimental support for the ego-dystonic nature of OCD and suggesting that habitual action is not sustained by dysfunctional belief.

## INTRODUCTION

Action is controlled by different learning mechanisms. On the one hand, actions followed by a reinforcer are more likely to be repeated in the future in a habitual fashion as a consequence of strengthening a stimulus-response representation. On the other hand, animals do not merely repeat previously reinforced actions but can instead make deliberate, goal-directed choices based on their knowledge of the relationship between an action and the associated outcome and their motivation to obtain that outcome [1]. As such, independent neural systems underlying goal-directed and habitual behavior orchestrate action control and such a delicate balance is essential for adaptive everyday behavior. Imbalance of the goal-directed and the habitual system has been hypothesized to be relevant for understanding compulsive behaviors [2] which manifest as actions persistently repeated without relationship to the overall goal [3]. Compulsions are also characterized by the feeling of being compelled or forced to engage in such behaviors [4] and they are generally associated with the insight that such actions are ultimately harmful and purposeless. Therefore, compulsive behaviors might be rooted in a disrupted synergy between the goal-directed and the habitual system whereby the habitual system seemingly overtakes response control and actions are divorced from their goals [2]. Obsessive-compulsive disorder (OCD) can be regarded as the prototypical disorder of compulsivity, which we used here as a benchmark to test this hypothesis. OCD manifests clinically as a lack of goal-directed control over repetitive, ritualistic actions and intrusive thoughts. OCD is ego-dystonic in nature as patients are generally able to recognize their compulsive behaviors and thoughts as disproportionate, excessive, and maladaptive [5]. Often, it is this ‘disconnection’ between the responses OCD patients find themselves making, as opposed to the responses they know to be rational, that causes so much distress [6].

Traditionally, cognitive theories posited dysfunctional beliefs as a major driver of OCD symptoms, to which cognitive treatments are targeted [7,8]. More recently, however, experimental evidence showing a tendency for OCD patients to display habitual behavior at the expense of goal-directed actions [9–11] has suggested that OCD is a disorder of habitual control. Such imbalance between hypothetical goal-directed and stimulus-response (S-R) habitual control over behavior has been shown by using the experimental manipulation of instructed outcome devaluation, i.e. changes in the *value* of the outcome previously associated with the action, as an experimental manipulation for detecting habit-based control. Excessive habits were thus expressed as an irrelevant maintenance of behavior, manifested as a lack of sensitivity to such a manipulation [9–11].

However, learning theory has established that goal-directed agents are also sensitive to the *causal* relation (i.e. contingency) between the response and the reward: if instrumental responding continues when such contingencies are degraded, it is assumed to be under habitual (S–R) control [1]. This manipulation of contingency-based instrumental responding has been tested across species and found to be mediated by fronto-striatal neural circuitry [12–20] implicated in OCD [21] and other disorders of compulsivity such as drug addiction [22] and binge-eating disorder [23]. As the causal action-outcome association is diminished, a reduction in behavioral responding is usually observed and, in humans, lower estimates of causal influence on the occurrence of the outcome are reported verbally via explicit causal judgments. Here, we developed a novel behavioral paradigm based on contingency degradation [1,16] to test the robustness of causal associations between actions and outcomes in OCD.

Importantly, with this experimental manipulation, we measured not only the rate of behavioral adjustment following changes in the causal action-outcome relationship, but also how subjects perceived that causal relationship. Therefore, we were able to test whether patients with OCD, compared to healthy volunteers, (i) showed goal-directed control by modulating their behavior in response to contingency degradation; (ii) accurately reported action-outcome knowledge of the causal relationship between response and associated reward; and crucially, (iii) differentially used action–outcome knowledge to guide their behavior. Therefore, our experimental manipulation enabled the testing of two competing hypotheses.

Compulsive behaviors (e.g. checking or rituals to prevent harm) may be interpreted as attempts to establish control. In this respect, compulsions might result either from an increased sense of responsibility [7] or, in contrast, as superstitious behaviors carried out either to regain a subjective sense of control or because contingencies are misperceived [24–26]. However, patients with OCD generally recognize their behavior as irrational, and hence exhibit a dichotomy between their behavior and their beliefs about the effectiveness of their actions. Therefore, a correspondence between inflated (or deflated) perceived contingencies and behavior would argue in favor of cognitive accounts for OCD, whereby compulsive behavior is guided by erroneous cognitive interpretation of environmental cues. In contrast, accurate detection of action–outcome contingencies in the face of behavioral insensitivity to contingency manipulation would provide support for a dissociation between an accurate cognitive appraisal of the environment and a failure to use this knowledge to guide behavior. The ego-dystonic nature of OCD, whereby the urge to perform an action is associated with the knowledge that the action is excessive or irrelevant would resonate with the latter scenario. Here, we test this prediction and show it to be valid. In addition, by using the contingency degradation intervention and avoiding verbally instructed devaluation procedures [9,11] it will be more feasible to make translational comparisons across species [17].

## RESULTS

### Contingency degradation

We used the experimental manipulation of contingency degradation to study detection of action-outcome contingencies in a sample of 27 OCD patients and 27 matched controls (**Table S1** and **Material and Methods**). Throughout the experimental session, the standard measure of contingency, ΔP, indexed the relationship between performing an action and obtaining an outcome. ΔP was the difference between two conditional probabilities: the probability of receiving an outcome upon performance of an action [probability of outcome given the action, P(O|A), i.e. the probability of response-contingent outcome] and the probability of receiving an outcome in the absence of that action [probability of outcome given the absence of an action, P(O|∼A), i.e. the probability of a non-contingent outcome], such that ΔP=P(O|A)–P(O|∼A) [27]. To degrade the contingency, once agents have learned to perform an action to receive a reward with a certain probability, a schedule of non-contingent outcome delivery is superimposed. By increasing the frequency of non-contingent outcomes, the overall contingency (i.e. the causal association between an action and its consequences) is degraded, hence reduced, or becomes negative. If guided by the goal-directed system, an agent should stop responding in face of contingency degradation (**Figure 1A-C**). Measures of interest include the overall relationship between actual and perceived contingency, and between contingency and behavior, but also specific contingency transitions in which P(O|∼A) increases without changes to P(O|A): this manipulation degrades instrumental contingency without affecting the contiguity of actions and outcomes that drives S–R habits, so is a specific test for excessive habitual responding.

**Figure 1.**
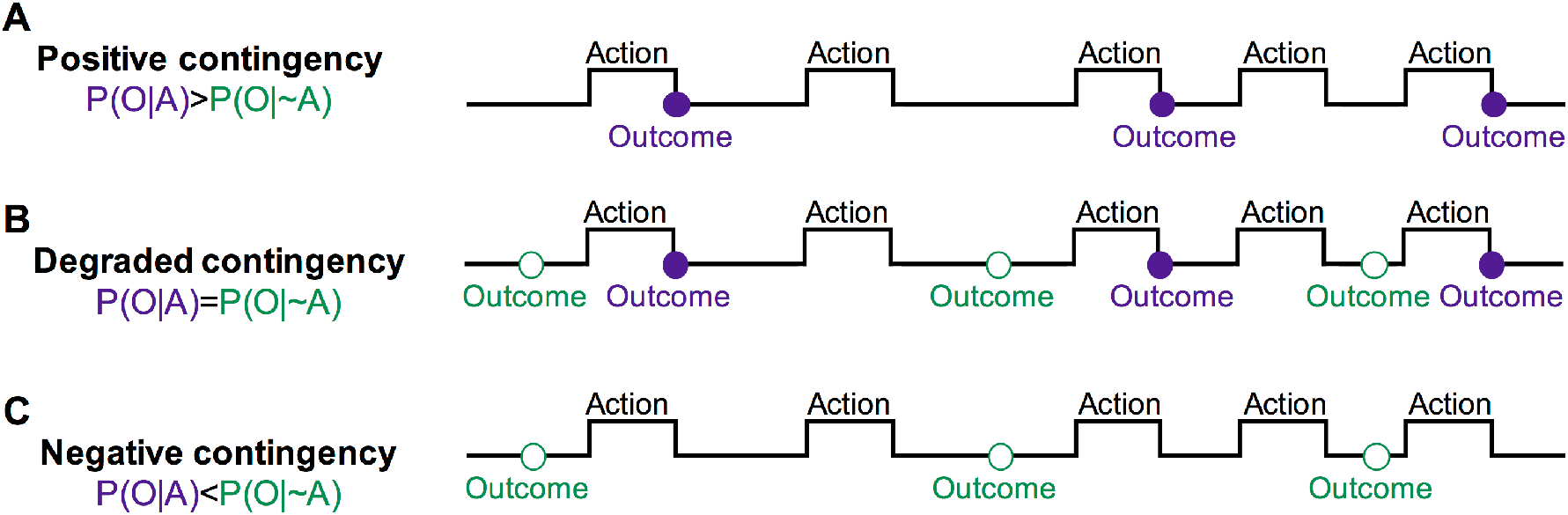
Contingency manipulation. **(A)** Diagram illustrating a schedule with a positive contingency, in which outcome is delivered upon performance of an action with a given probability P(O|A). **(B)** Contingency is degraded by also delivering outcomes in the absence of an action, with a given probability P(O|∼A). If the contingency is degraded to the extent that the two probabilities are equal, the causal status of the action is nil and the probability of the reinforcer is the same regardless of any response. **(C)** When the P(O|∼A) is higher than P(O|A), the contingency becomes negative and the action reduces the probability of reinforcer delivery. P(O|A), probability of outcome given the action, i.e. the probability of receiving a response-contingent outcome; P(O|∼A) probability of outcome given the absence of an action, i.e. the probability of a non-contingent outcome. Violet, filled circle for contingent outcomes; green, empty circle for non-contingent outcomes.

### A novel protocol to test sensitivity to action-outcome contingency

We developed and implemented a novel free-operant, self-paced procedure. The instructions informed the participants that they could earn 25 pence (p; £0.25) whilst pressing the space bar on a keyboard, and that they were free to press the key as often as they liked (**Figure 2 A** and **Material and Methods**). They were further instructed that the relationship between pressing the space bar and receiving the 25p reward would vary during the experiment, and that pressing the space bar might earn a reward, a reward might also arrive on its own, or pressing the space bar might prevent a reward from arriving. Lastly, they were informed that occasionally they would be asked to rate the degree to which pressing the space bar caused the occurrence of the reward. We varied P(O|A) and P(O|∼A) to give blocks with different levels of contingency and obtain different experimental conditions (**Figure 2 B, C** and **Table 1**). In positive contingency conditions, P(O|A) was higher than P(O|∼A). Those were degraded by increasing P(O|∼A). To mimic the maladaptive nature of compulsivity in OCD, by which actions are repeated despite adverse consequences, negative contingencies were also introduced in the experimental paradigm whereby P(O|∼A) was higher than P(O|A). In these situations, performing the action reduced the probability of getting an outcome.

**Figure 2.**
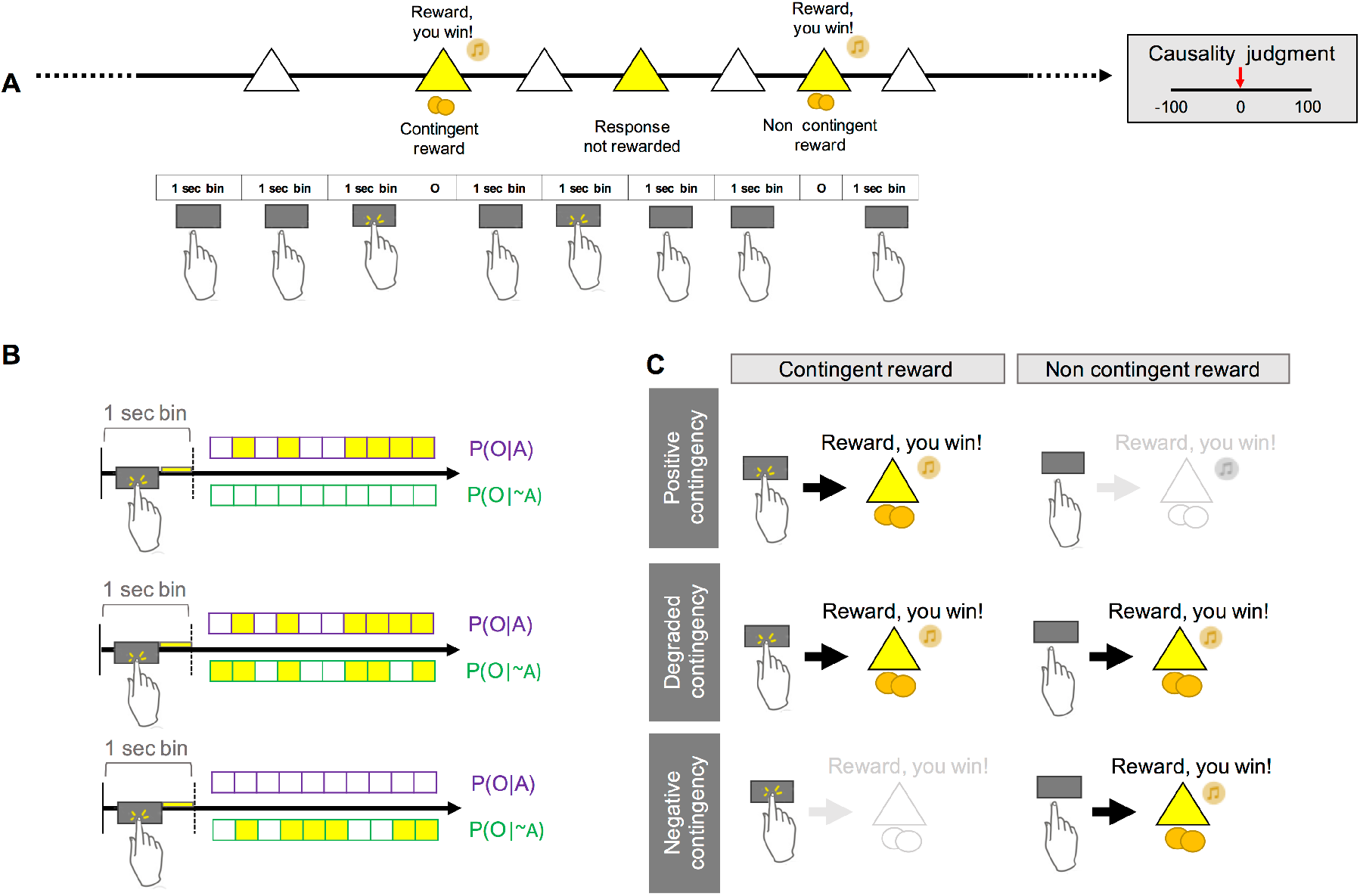
Experimental paradigm. **(A)** Subjects had to complete an experimental session of 12 blocks of 2 minutes each. At the end of each block, subjects had to judge to what extent pressing the space bar caused the occurrence of the reward, on a scale from −100 (pressing the space bar always prevented reward) to 100 (pressing the space bar always caused reward). During the experimental session, the participant was presented with a white triangle and could decide whether to press the space bar or not. Rewards were delivered contingently upon pressing of the space bar or non-contingently in the absence of a response. In addition, a running total of the amount of money earned within a block was continuously displayed in the upper corner of the screen (not shown in figure). Note that in cases where the participant was not pressing the space bar for multiple (hidden) 1 sec bins in a row, the white triangle was continuously displayed on the screen, unless a non-response contingent reward occurred. In those cases, a reward was displayed on the screen non-contingently. **(B)** Each block was divided into 120 unsignaled time periods (bins) of 1 second. When a response occurred within each bin, the triangle turned yellow until the bin ended. If a response was recorded during the bin, a contingent reward was delivered at the end of that bin according to the applicable probability of outcome delivery given a response, P(O|A). If no response occurred during the bin, a non-contingent reward was delivered according to the applicable probability of outcome delivery given the absence of a response, P(O|∼A). **(C)** By varying P(O|A) and P(O|∼A), different levels of contingencies were achieved so that each experimental session included positive, degraded, and negative contingency blocks. P(O|A), probability of outcome given the action, i.e. probability of receiving a response-contingent outcome; P(O|∼A) probability of outcome given the absence of an action, i.e. probability of a non-contingent outcome.

**Table 1.** Response rates and causality judgments.

|  |  | Programmed |  |  | Experienced |  | Response |  | Causality |  |
| --- | --- | --- | --- | --- | --- | --- | --- | --- | --- | --- |
|  |  | contingency |  |  | contingency |  | rate |  | judgment |  |
| Block |  | P(O\|A) | P(O\|~A) | ΔP | CTL | OCD | CTL | OCD | CTL | OCD |
| Fixed Order | 1 | 0.60 | 0.00 | 0.60 | 0.59 | 0.60 | 0.51<br>(0.21) | 0.49<br>(0.24) | 43.30<br>(34.27) | 48.60<br>(31.82) |
|  | 2 | 0.60 | 0.60 | 0.00 | 0.01 | 0.05 | 0.26<br>(0.20) | 0.35<br>(0.27) | 8.17<br>(27.64) | 10.67<br>(44.74) |
|  | 3 | 0.00 | 0.00 | 0.00 | 0.00 | 0.00 | 0.37<br>(0.27) | 0.48<br>(0.27) | -10.46<br>(39.75) | -14.81<br>(45.60) |
| Shuffled in a Latin square design | 4 | 0.00 | 0.00 | 0.00 | 0.00 | 0.00 | 0.38<br>(0.23) | 0.48<br>(0.27) | -15.17<br>(36.06) | -21.35<br>(40.43) |
|  | 5 | 0.00 | 0.30 | -0.30 | -0.29 | -0.30 | 0.26<br>(0.21) | 0.27<br>(0.21) | -50.43<br>(47.60) | -41.51<br>(55.30) |
|  | 6 | 0.00 | 0.60 | -0.60 | -0.62 | -0.60 | 0.20<br>(0.23) | 0.21<br>(0.16) | -55.27<br>(44.83) | -32.53<br>(66.08) |
|  | 7 | 0.30 | 0.00 | 0.30 | 0.30 | 0.30 | 0.49<br>(0.22) | 0.62<br>(0.20) | 27.54<br>(24.20) | 35.34<br>(25.33) |
|  | 8 | 0.30 | 0.30 | 0.00 | 0.03 | 0.00 | 0.34<br>(0.26) | 0.41<br>(0.24) | 0.01<br>(31.53) | 0.36<br>(37.34) |
|  | 9 | 0.30 | 0.60 | -0.30 | -0.28 | -0.29 | 0.29<br>(0.26) | 0.32<br>(0.26) | -12.95<br>(47.98) | -9.40<br>(38.56) |
|  | 10 | 0.60 | 0.00 | 0.60 | 0.60 | 0.60 | 0.62<br>(0.21) | 0.56<br>(0.23) | 56.01<br>(26.67) | 53.52<br>(30.84) |
|  | 11 | 0.60 | 0.30 | 0.30 | 0.32 | 0.31 | 0.38<br>(0.26) | 0.53<br>(0.25) | 22.49<br>(30.71) | 33.06<br>(32.79) |
|  | 12 | 0.60 | 0.60 | 0.00 | -0.01 | 0.00 | 0.29<br>(0.23) | 0.38<br>(0.25) | 9.64<br>(31.81) | 8.30<br>(36.75) |

$P(O|A)$ , probability of the outcome given the action; $P(O|\sim A)$ , probability of the outcome in the absence of the action; $\Delta P$ =contingency. Dependent variables are given as mean (SD).

### Effect of instrumental contingency on response rate

In line with previous data in healthy volunteers, mean response rate increased with contingency (contingency, F_4,208_ = 65.028, p < 0.001) (**Table 1** and **Figure 3 A**). Overall levels of responding did not differ between the groups (group, F_1,52_ = 1.074, p = 0.305), ruling out apathy or, in contrast, generalized impulsivity, in the OCD group. Responding in the groups was differentially affected by the contingency (group×contingency, F_4,208_ = 3.922, p = 0.01); this difference was explored via between-groups simple-effect comparisons at each level of contingency. Patients with OCD persisted in responding more than healthy subjects in the face of reduced instrumental contingency (group ΔP = 0.3, F_1,52_ = 6.036, p = 0.017) (specific transitions in which P(O|∼A) increased without changing P(O|A) are explored further below). Increased response rates in patients did not correlate with impulsivity traits, measured by the Barratt Impulsiveness Scale [28] (r = 0.312, p = 0.129). Patients responded marginally more at ΔP = 0.0, but this did not reach significance (F_1,52_ = 3.185, p = 0.080).

**Figure 3.**
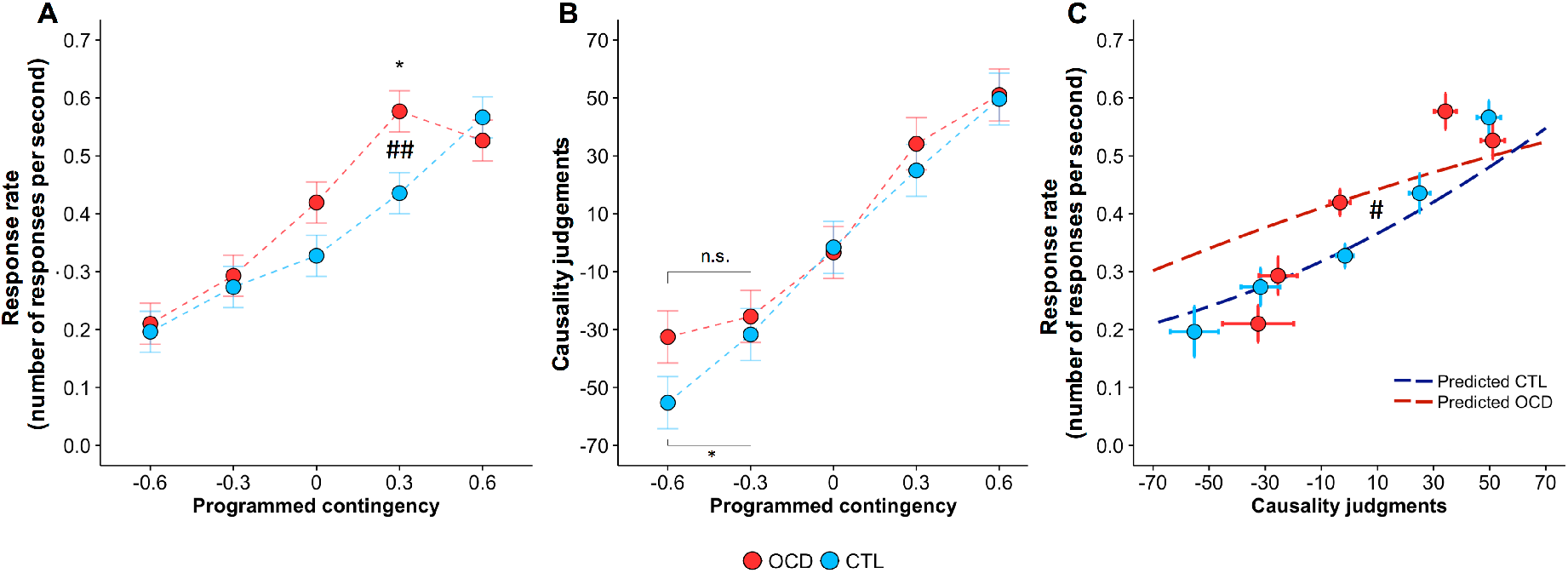
Increased response rate but intact action-outcome knowledge and their dissociation in OCD. (A)Mean response rate by contingency. Both groups responded more for higher contingencies. However, OCD patients showed reduced sensitivity to instrumental contingency. ##p < 0.01, interaction; *p < 0.05, for between-group comparison. **(B)** Subjective judgments of causality increased as a direct function of response–outcome contingency in both groups. Data are presented in ascending order of programmed contingency, but contingencies were experienced by each subject in a semi-randomized order. Error bar indicates Fisher’s Least Significant Difference (FLSD) to facilitate post-hoc comparisons (error bars are ± 0.5 × tcritical × SD). However, in the context of mixed designs, as in this case, this error bar can only be used for within-subject comparisons. The difference between OCD and CTL in mean causality judgments at ΔP = −0.6 was not significant. However, controls but not OCD patients subjectively detected a difference between neighboring levels of negative programmed contingency between ΔP = −0.3 and ΔP = −0.6). *p < 0.05, for within-group comparison. **(C)** Response rate as a function of causality judgment. The two groups differentially employed action–outcome knowledge to guide their behavior (# p < 0.05, group × quadratic causality judgment interaction). Dashed lines show predictions from the best-fit model (predicting response rate using group and both quadratic and linear effects of causality judgments); points/error bars (SEMs) show values clustered by programmed contingency. The apparent discrepancy for strongly negative causality judgments reflects the fact that the model uses within-subject regression and that not all patients gave causality ratings that extended to the left-hand end of the range (see **Figure S2**). CTL, controls; OCD, patients with obsessive–compulsive disorder. Programmed contingency refers to the a priori experimentally programmed contingency, resulting from the a priori programmed conditional probabilities. As described in the main text, data were collapsed across blocks having equal contingencies [ΔP = - 0.6, Block 6; ΔP = - 0.3, Block 5, Block 9; ΔP = 0.0, Block 2, Block 3, Block 4, Block 8, Block 12; ΔP = 0.3, Block 7, Block 11; ΔP = 0.6, Block 1, Block 10. See Table 1 for naming of the blocks]; specific contingency transitions to detect habitual responding are shown inFigure 4.

The group difference in the effect of contingency remained significant even when considering only medicated OCD and controls (group _OCD medicated, Controls_ ×contingency, F_4,176_ = 4.107, p = 0.003) or only unmedicated OCD and controls (group _OCD unmedicated, Controls_ ×contingency, F_4,132_ = 2.628, p = 0.037). There were no between-group effects nor interactions that depended on medication status in OCD patients (all p > 0.1) (**Figure S1**).

We recorded the number of presses made within each 1-s time bin and did not detect a difference between groups in the number of additional number of responses within each bin (those beyond the first such response, which had behavioural effects). ‘Additional’ (superfluous) responding was not affected by instrumental contingency (contingency, F_4,208_ = 0.621, p = 0.648) or group (group, F_1,52_ = 0.017, p = 0.896; group×contingency, F_4,208_ = 0.070, p = 0.991). Differences in the additional number of responses within each bin would have been consistent with a framework in which excessive responding in OCD is attributed to a failure of inhibition. Our findings instead reinforce the notion that OCD patients expressed habitual responding, a hypothesis we test directly below.

### Effect of instrumental contingency on causality judgments

Causality ratings were a function of action-outcome contingency across both groups (**Table 1** and **Figure 3B**) (contingency, F_4,208_ = 74.099, p < 0.001). The two groups did not differ in their judgements of causality (group, F_1,52_ = 2.379, p = 0.129; group×contingency, F_4,208_ = 1.084, p = 0.366). The results did not change when considering only medicated OCD and controls or only unmedicated OCD and controls. There were no between-group effects nor interactions that depended on medication status in OCD patients (all p > 0.186) (**Figure S1**).

### Relationship between response rate and causality judgments

Patients with OCD and controls differed in the way that causality judgements predicted response rate, in a non-linear fashion. Overall, response rate was linearly predicted by causality ratings (F_1,45.449_ = 58.154, p < 0.001). We did not identify a difference in this linear relationship between groups (group x causalitylinear: F_1, 45.449_ = 1.489, p = 0.229). However, there was a significant non-linear effect as well, which differed between groups (group×causalityquadratic

F_1,204.827_ = 3.959, p = 0.0479) (**Figure 3C**). Residuals were larger in the OCD group (F test of residual variances by group: F_323,323_ = 1.28, p = 0.013), indicating a slightly poorer model fit in OCD; however, the residual variance was only 28% larger (controls 0.024; OCD 0.0308) which does not jeopardize the group comparisons [29].

This indicated an altered, non-linear relationship between causality judgments and response rate in patients and represents a formal demonstration of the differential and blunted use of action-outcome knowledge to modulate behavior in patients, also supported by patients’ reports (**Table 2**). Thus, in patients, for positive contingencies, behavior persisted after contingency degradation despite intact and accurately reported action-outcome knowledge of the causal effect of their actions. For negative contingencies, the best-fit model predicted increased response rate in patients when they believed their actions to be detrimental. The equal response rates (**Figure 3A**) may have been a consequence of this effect plus a non-significant tendency to believe their actions to be less detrimental than controls at highly negative contingencies (**Figure 3B**, programmed contingency –0.6). We analyzed response rate for different time windows of each block, excluding the possibility that such dissociation was due to different learning processes in OCD patients (**Figure S3**). Habitual responding emerged towards the end of each block, closer in time to when subjects reported their causality judgments. This rules out the possibility that OCD patients were simply slower to learn the contingency: habitual responding was observed at times close to subjective causality judgments for which OCD patients did not differ from controls.

**Table 2.**
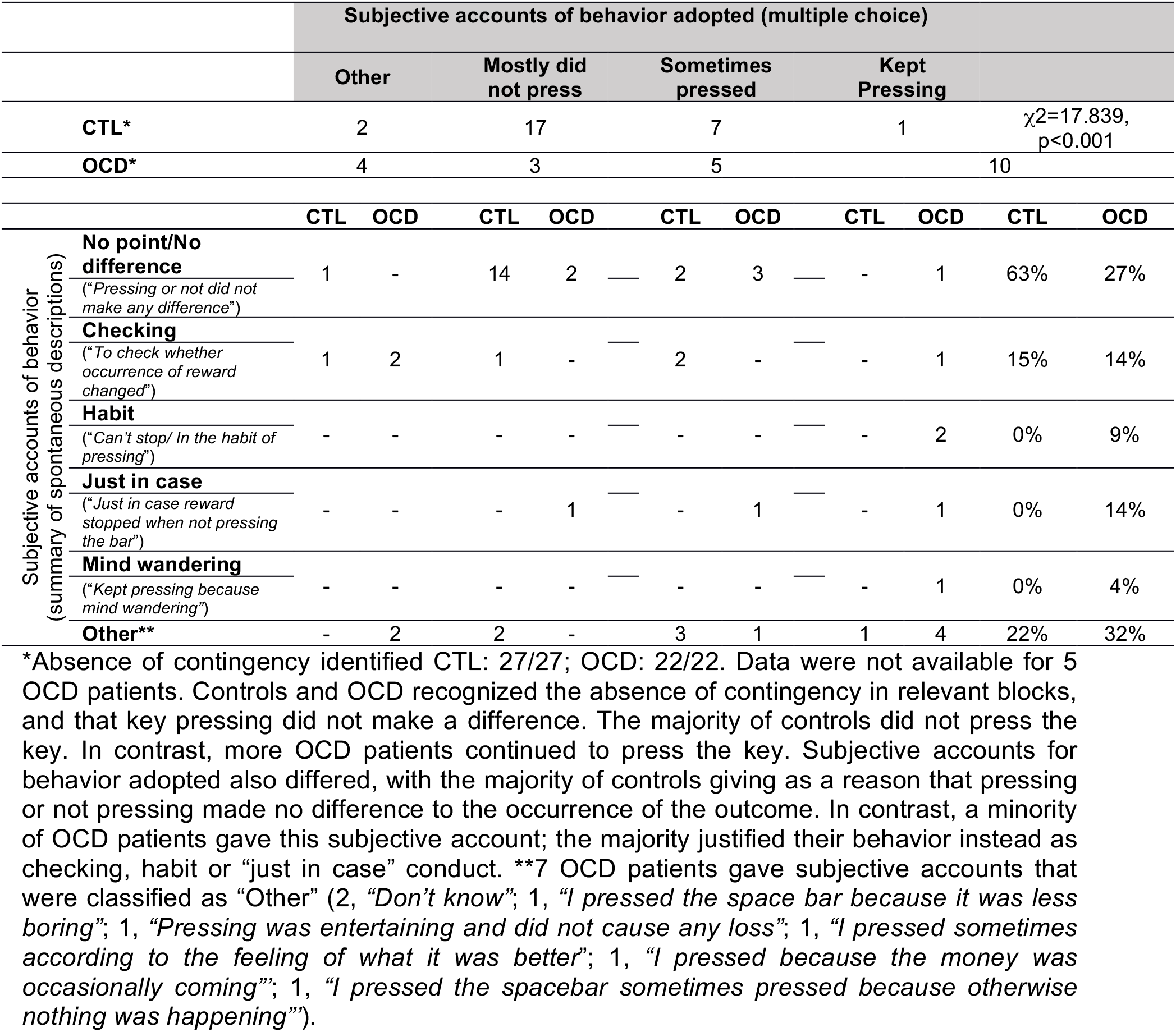
Subjective accounts when the contingency was zero.

### Habit/goal-directed ratio score

We tested for differences in habitual responding directly, by examining contingency transitions in which P(O|A) was positive and held constant and P(O|∼A) was increased, to test precisely if increased responding observed for ΔP = 0.3 (**Figure 3A**) was due to habitual behavior. To match number of observations for each condition, we focused on contingency degradation occurring after the implicit training phase. We therefore compared responding for blocks in which P(O|A) was held constant at 0.6 and P(O|∼A) increased from 0 to 0.3 leading to a degraded contingency of ΔP = 0.3 (Block 10, Block 11), by computing a ratio score. On this measure (contingent/(contingent+degraded see Material and Methods), which controls for response variability across subjects, high scores (close to 1) indicate responsivity to the contingency change, and low scores (close to 0.5) indicate habitual responding. Whereas control subjects showed a robust decline in responding upon contingency degradation, as indicated by a ratio-score well above 0.5 (one-sample t test tested against 0.5, t_26_ = 5.918, p < 0.001) patients with OCD responded nearly equally in both conditions, with their ratio-score being close to 0.5 (one-sample t test tested against 0.5, t_26_ = 0.585, p = 0.563). There was a significant between-groups difference in the ratio-score (t_52_ = 3.350, p = 0.002) (**Figure 4A**). Furthermore, subjects were classified dichotomously as ‘goal-directed’ (ratio-score>0.5) or ‘habitual’ (ratio-score≤0.5). A higher proportion of ‘habitual’ subjects was found in the OCD group (controls habitual 2/27; OCD habitual 12/27; χ^2^_1_ = 7.811, p = 0.005). There was no correlation between the ratio-score and symptom severity (Y-BOCS) in OCD patients (r = −0.101, p = 0.625).

Similarly, we observed a marginal effect for increased responding when ΔP = 0.0 (**Figure 3A**). Therefore, we calculated a ratio-score for blocks for which the action-outcome relationship was contingent (ΔP = 0.6, P(O|A) = 0.6, P(O|∼A) = 0.0, Block 10) and then completely degraded to ΔP = 0.0 by superimposing a non-contingent schedule (ΔP = 0.0, P(O|A) = 0.6, P(O|∼A) = 0.6, Block 12). Even though both groups showed ratio scores significantly different from 0.5 (one-sample t test tested against 0.5, controls, t_26_ = 7.334, p < 0.001; OCD, t_26_ = 3.388, p = 0.002), OCD patients showed diminished goal-directed behavior compared with controls (ratio-score, t_52_ = 2.23, p = 0.03) (**Figure 4B**). There was no difference between OCD and controls in the ratio score for when action-outcome relationship was contingent (ΔP = 0.3, P(O|A) = 0.3, P(O|∼A) = 0.0, Block 7) and then completely degraded to ΔP = 0.0 by superimposing a non-contingent schedule (ΔP = 0.0, P(O|A) = 0.3, P(O|∼A) = 0.3, Block 8) (**Figure 4C**).

**Figure 4.**
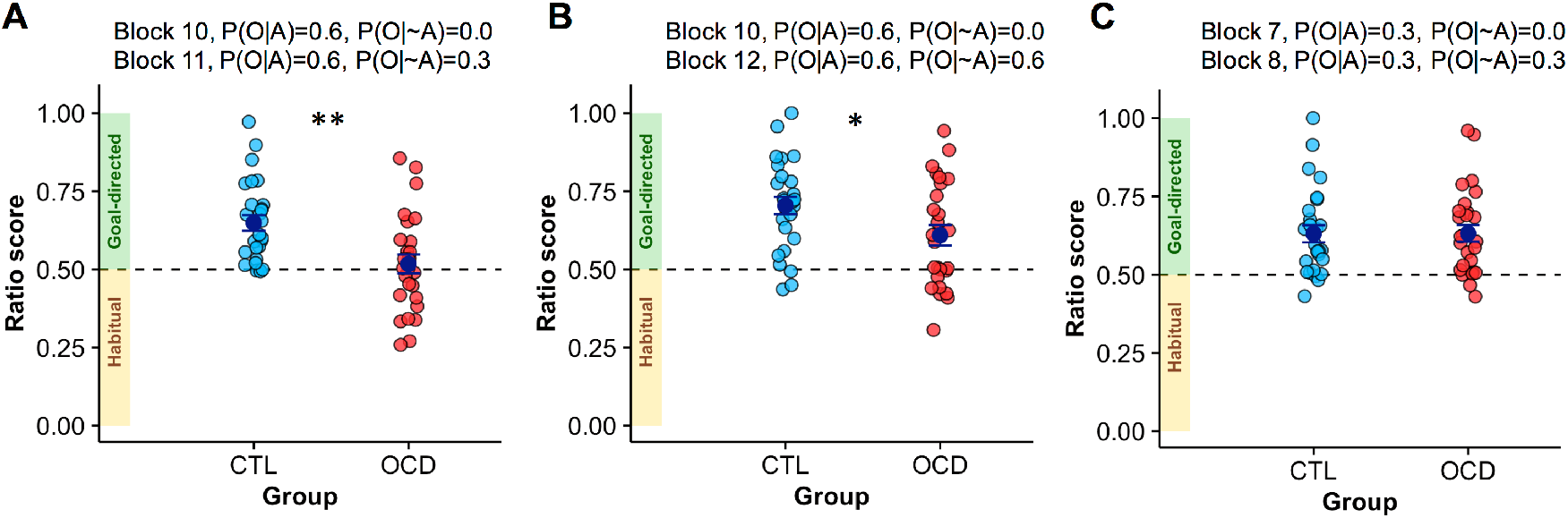
Habit/goal-directed ratio-scores for contingent and corresponding degraded-contingency conditions identifies habitual responding in OCD patients. A ratio score was calculated for pairs of blocks across which the contingency was degraded by keeping P(O|A) constant and increasing P(O|∼A). The first block of each pair is termed “contingent”, as a shorthand, and the second “degraded”; the ratio score was then calculated as contingent/(contingent+degraded). High scores (close to 1) indicate that the subject responds to the degradation; low scores (close to 0.5) indicate insensitivity to the degradation and therefore habitual responding. **(A)** Ratio scores for pairs of blocks for which the contingency was degraded from ΔP = 0.6 (P(O|A) = 0.6, P(O|∼A) = 0.0, Block 10) to ΔP = 0.3 (P(O|A) = 0.6, P(O|∼A) = 0.3, Block 11). OCD patients displayed increased habitual behavior (t_52_ = 3.350, p = 0.002). **(B)** Ratio scores for pairs of blocks for which the contingency was degraded from ΔP = 0.6 (P(O|A) = 0.6, P(O|∼A) = 0.0, Block 10) to ΔP = 0.0 (P(O|A) = 0.6, P(O|∼A) = 0.6, Block 12). OCD patients showed increased habitual behavior compared with controls (t_52_ = 2.23, p = 0.03). **(C)** Ratio scores for pairs of blocks for which the contingency was degraded from ΔP = 0.3 (P(O|A) = 0.3, P(O|∼A) = 0.0, Block 7) to ΔP = 0.0 (P(O|A) = 0.3, P(O|∼A) = 0.3, Block 8). Error bars: SEM. CTL, controls; OCD, patients with obsessive– compulsive disorder. *p < 0.05; **p < 0.01.

The response to contingency degradation was impaired in OCD patients when degradation occurred from high baseline contingency (ΔP = 0.6, **Figure 4A-B**) but not from low contingency (ΔP = 0.3, **Figure 4C**). We therefore investigated responses rate at high and low instrumental contingencies in OCD patients and controls. OCD patients responded more than controls when the overall instrumental contingency was low (ΔP = 0.3, P(O|A) = 0.3, P(O|∼A) = 0.0, Block 7) (OCD = 0.62 ±0.20; Controls = 0.49 ±0.22; post hoc t-test F_52_ = 4.961, p = 0.030, **Table 1** and **Figure S4**). In addition, OCD patients did not show significant modulation of response rate from high (ΔP = 0.6 [P(O|A) = 0.6, P(O|~A) = 0.0], Block 10) to low (ΔP = 0.3 [P(O|A) = 0.3, P(O|-A) = 0.0], Block 7) instrumental contingency (Patients, Block 10: 0.56±0.23; Patients, Block 7: 0.62±0.20). In contrast, controls did show such modulation (Controls, Block 10: 0.62±0.21; Controls, Block 7: 0.49 ±0.22). There was in fact a significant interaction (F_1,52_ = 11.674, p = 0.001) between Group (Control, Patients) and Block (Block 7, Block 10). These findings therefore suggest OCD patients had increased response rate when there was a low instrumental contingency between the action and the outcome, although they were able to modulate their response rate when a contingency degradation occurred against the background of such low contingency.

To test the effect of repetition in the development of habits, we computed the ratio score for the early phases of the experimental design (Early: Block 1 and Block 2) and compared with late ones (Late: Block 10 and Block 12). There was no main effect of time (F_1,52_ = 0.083, p = 0.775) nor a time × group interaction (F_1,52_ = 0.648, p = 0.425). Therefore, we did not detect an effect of repetition in the development of habits [30]. Across groups, habitual behavior in the early phases of the experimental design was associated with higher OCD traits measured by the OCI-R (r = −0.280, p = 0.046).

### Absence of depressive realism in OCD

Previous data have shown that healthy non-depressed subjects have biased higher estimates of causality judgments when the contingency is zero [31]. This erroneous estimation arises when contingent and non-contingent outcomes occur frequently (i.e. high density of reinforcement), but not when contingent and non-contingent outcomes occur infrequently (i.e. low density of reinforcement). In contrast, depressed individuals show a “depressive realism” whereby, irrespective of the density of reinforcement, correctly report having no causal effect on the occurrence of the outcome [31]. Because OCD patients showed higher depression scores compared with healthy subjects, we tested possible between-group differences in causality judgments for ΔP = 0.0 blocks with different densities of reinforcement (Block 4, 8, and 12, **Table 1**). Selection was limited to the Latin square phase to have an equal number of observations for each condition. Estimation of control was higher for higher reinforcement density (F_2,104_ = 8.365, p < 0.001) (**Figure 5A**), with no between-groups differences (group, F_1,52_ = 0.171, p = 0.681; group×reward density, F_2,104_ = 0.124, p = 0.883), despite higher levels of depressive symptoms in OCD patients.

**Figure 5.**
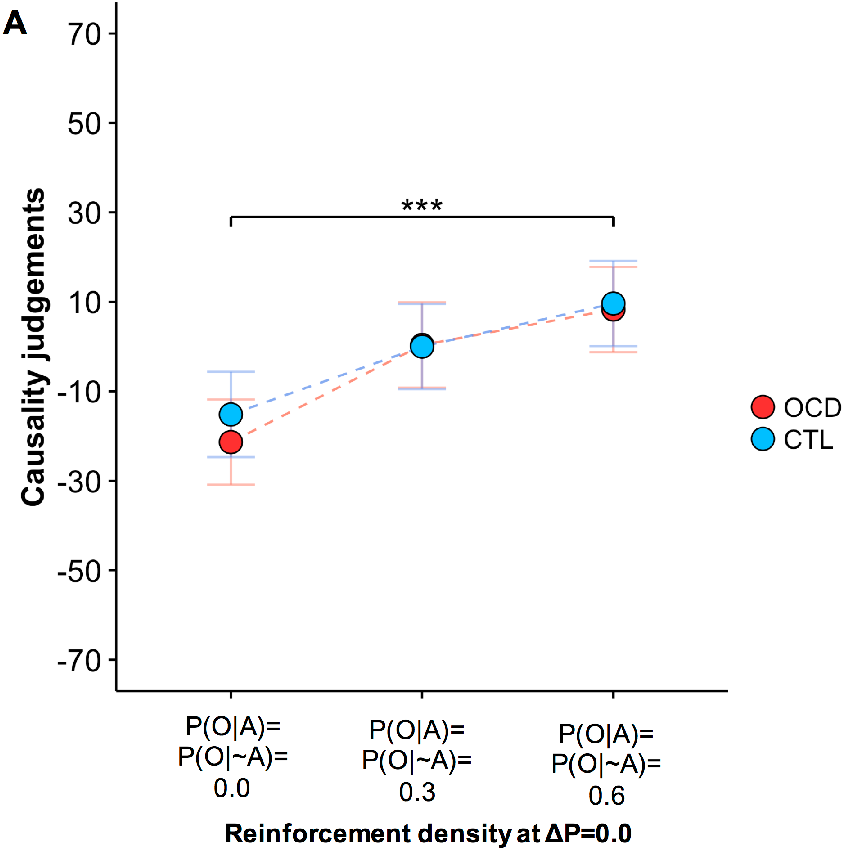
OCD patients show intact causality judgments when when the contingency was zero proving absence of ‘depressive realism’. **(A)** For both controls subjects and patients with OCD, causality judgments increased as a function of higher density of reinforcement even though there was no causal association between the action and the outcome (contingency ΔP = 0.0) in all three situations. Error bar indicates Fisher’s Least Significant Difference (FLSD) to facilitate post-hoc comparisons (error bars are ± 0.5 × t_critical_ × SD). However, in the context of mixed designs, as in this case, this error bar can only be used for within-subject comparisons. ***p < 0.001, main effect of density of outcome. CTL, controls; OCD, patients with obsessive–compulsive disorder.

## DISCUSSION

Our findings show a failure in a learning process regulating action control based on the relationship between actions and their consequences in a sample of individuals characterized by clinically relevant high levels of compulsive behaviors. Patients with OCD exhibited increased response rates when outcomes were less contingent upon responding (**Figure 3**), very likely as a consequence of enhanced S-R habitual tendencies (**Figure 4**). In contrast, explicit action–outcome knowledge was intact: patients were capable of accurate subjective assessments of the cause–effect relationship between actions and their consequences, which did not differ from those of controls. However, in patients, action– outcome knowledge did not translate normally into action. Increased habitual behavior was dissociated from intact explicit action–outcome knowledge about the effectiveness of their actions. Moreover, in patients, response rate was augmented when they believed their actions to be detrimental (**Figure 3C**).

OCD patients exhibited excessive, presumed habitual, responding when the action–outcome contingency was degraded, and the action was thus less causally linked to an outcome, the effect being present when contingency was partially and completely degraded. The relationship between subjective causality judgement and behaviour also predicted excessive responding for actions patients believed to be detrimental (**Figure 3C**), in keeping with the clinical manifestation of persistent behavior even when recognized as being harmful.

Our results showed that OCD patients showed habitual behavior when degradation occurred from high levels of contingency (4A, 4B and Supplementary S4A, S4B). When the degradation occurred from low levels of contingency, OCD patients were not habitual (4C and Supplementary S4C). While no *a priori* hypothesis was formulated, we observed that OCD patients kept their response rate constant regardless of whether there was high or low contingency between the action and the outcome. This behavior highlights how OCD patients exhibited increased response rates when there was a low instrumental contingency between the action and the outcome.

Translational work in rats [12], marmoset monkeys [17], and humans [18,20] using contingency degradation has shed light on possible cortico-striatal determinants of goal-directed and habitual actions. In rats, pharmacological manipulation of the prelimbic cortex and the dorsomedial striatum (the putative homologue of the caudate nucleus in humans) prevented the encoding of action–outcome associations during instrumental conditioning [32]. In marmosets, insensitivity to contingency degradation has been found following lesions to the perigenual anterior cingulate cortex and the orbitofrontal cortex [17]. In humans, activity in the medial prefrontal cortex (PFC)/medial orbital cortex and the anterior caudate is associated with contingency learning and goal-directed behavior [18,20,33]. In healthy volunteers, reduced gray matter volume in the caudate correlates with a propensity towards habits [23]. In OCD patients, hyperactivity of the caudate nucleus is associated with excessive habit formation, tested in avoidance by means of outcome devaluation [10]. These fronto-striatal regions are implicated in the pathophysiology of OCD [21]; therefore, habitual responding, manifested as lack of behavioral suppression of action upon contingency degradation, plausibly depends on abnormalities in such circuits. Animal work has also shown differential sensitivity to outcome devaluation and contingency degradation [34]. Such a distinction can be now tested in humans as well, using the paradigm devised here and those focusing on outcome devaluation. By contrast, response rates in control participants faithfully tracked the level of instrumental contingency, in line with other studies in healthy populations [18–20].

Accurate subjective judgments in both groups on the causal relationship between actions and outcomes, especially in the case of positive contingencies, indicated intact action– outcome knowledge not only in controls, as previously shown [13,18–20], but also in OCD. For OCD patients, we also observed that for negative contingencies only, actions were reported to be less detrimental than experienced. Although these findings should be interpreted with caution in the context of a lack of a main effect, they might relate to the maladaptive nature of OCD where actions are repeated despite negative consequences. Here, our findings might suggest slight inaccuracies of subjective judgement in case of negative contingencies which might contribute to patients’ perception of their actions to have less disadvantageous consequences than experienced.

Previous studies have shown that affective states influence how objective contingencies are perceived [31]. In situations in which there is a lack of action–outcome contingency, overestimation of causal control is observed in non-depressed people when the non-contingent reward occurs frequently. Such an effect is not found in depressed individuals, who show an accurate detection of the lack of contingency (i.e. “depressive realism”). In the present study, when the contingency was zero, causality judgments increased as a direct function of the density of the reward and equally in controls and patients.

In addition, even if patients were relatively more depressed than healthy volunteers, their emotional/affective state did not influence their perception of the environmental contingencies in a way that was significantly different from that of healthy volunteers (**Figure 2B**).

Our findings also demonstrate that OCD patients used their knowledge about environmental contingencies to guide their actions in a manner that differed from controls. Subjective detection of instrumental contingency was dissociated from expressed behavior in patients. For positive contingencies [P(O|A) > P(O|∼A)], OCD patients displayed increased response rates but accurate subjective reports of contingency. For negative contingencies, [P(O|A) < P(O|∼A)], in OCD patients response rates were not affected but there was evidence of contingencies being reported as less detrimental than experienced. Increased response rates were observed when the introduction of non-contingent outcomes reduced contingency, and inaccurate contingency ratings were observed when the non-contingent outcomes were more likely than contingent ones. Even though it remains to be clarified why non-contingent outcomes had a differential effect on behavior and reported causality judgments, it appears plausible that patients had particular difficulties in integrating non-contingent conditional probabilities. Such an effect might be dependent on a circuit including the posterior caudate and the inferior frontal gyrus, which has been shown to selectively decode non-contingent conditional probabilities [18].

Previous studies have shown that functional activity of the inferior and superior parietal lobule and the middle frontal gyrus scales with subjective reports of instrumental contingency [18]. Parietal abnormalities [35] together with diminished caudate–parietal connectivity [36,37] characterize OCD. Such abnormalities might contribute to inefficient use of explicit knowledge of instrumental contingencies to guide behavior in OCD. Therefore, these observations prompt the hypothesis that the inability to modulate behavior according to action–outcome contingencies in OCD patients might be due not only to abnormal striatal encoding of action–outcome contingencies, but also (or alternatively) to an inability of action–outcome metacognitive knowledge (putatively dependent on parietal activity) to guide behavior. In this respect, empirical testing will clarify if a lack of integration between the fronto-parietal system and the caudate nucleus contributes to the ego-dystonic, compulsive nature of OCD.

More generally, cognitive theories of OCD [7,38] conceptualize the disorder in light of an exaggerated appraisal of intrusive thoughts, which is believed to be the critical factor in the maintenance of the disorder. In this respect, OCD is identified in terms of the impact of inflated evaluation of intrusive negative thoughts on action. In the present study, in direct contrast, patients with OCD showed intact knowledge of the contingency, especially in the case of positive contingencies, between the action and the outcome but exaggerated responding despite this correct appraisal of contingencies. Therefore, even if OCD subjects were aware of the contingency, they did not use it to guide their behavior. Rather than supporting a model whereby OCD is maintained by exaggerated and dysfunctional appraisal of action contingencies, the findings suggest that exaggerated actions, possibly rooted in a propensity towards the development of habits [39], lie at the core of the disorder.

We found increased behavioral reliance on habits in OCD, in agreement with previous studies which have shown habitual behavior in OCD by using outcome devaluation in appetitive [9] and aversive domains [11]. Here, we have extended those findings by testing habits via contingency degradation as defined by Dickinson and colleagues [1].

We found a correct appraisal of the contingency between action and outcome, in line with previous data showing intact awareness of explicit associative contingencies in case of outcome devaluation in this patient population [11], though in a context of multiple action-outcomes associations OCD patients show weaker knowledge on the causal relationship between actions and their respective outcomes [9]. Imbalances between the goal-directed and the habitual system in OCD, which we identified here, has also been shown in OCD using neurocomputational models [23].

Finally, our work has theoretical implications for understanding goal-directed and habitual systems. In fact, it is common to assume competition between these two systems (i.e., that if a behavior is not under goal control, then it must be a habit). However, this study contributed to the relevant literature in showing that goal-directed and habitual forms of behavior can coexist in accordance with recent views [40]. Namely subjective reports in OCD patients tracked goal-directed contingencies correctly, while behavior was presumably habitual. Therefore, this evidence suggests that adaptive behavior depends on a fine tuning and coordination between the two systems, which probably go awry in OCD patients.

By contrast with classical theories predicting development of habits due to repetition over time [30], we did not observe a shift from goal-directed to habitual behavior over early and late experimental phases. There was no statistical effect of repetition. This might be due to a limitation of the experimental design, which did not lend itself to an optimal investigation of this aspect. In fact, participants experienced the first three blocks in the same order at the beginning of the experiment, but blocks were then presented in a semi-randomized design. This manipulation might conceivably have diluted the effect of repetition due to the different number of instrumental contingency blocks experienced prior to the relevant critical test across subjects. In addition, the relatively short duration of the task may have limited the possibility of detecting such training effects. Recent evidence also suggests that limited overtraining in instrumental behaviors fails to enhance development of habits (de Wit et al., submitted). In the initial phase of the experimental session, there was variability in the extent to which goal-directed or habitual strategies were adopted in both groups and a tendency for an association with OCD traits. However, as this correlation was not marked and was observed only when considering the whole sample, replication is warranted. OCD is known to be linked to abnormalities of serotonergic function, and there is evidence in healthy humans that diminished serotonin neurotransmission promotes habitual behavior [41]. We did not find an effect of medication, but given the small samples size we had insufficient power to draw definite conclusions. However, it seems unlikely that the effect observed was due to medication status of the OCD patients, as such medication is designed to increase serotonergic transmission.

In conclusion, this study reinforces the hypothesis that habit formation is a contributor to a disorder of compulsivity, using a novel, independent, valid behavioral assay based on contingency degradation that can readily be translated across species. A mismatch between explicit action–outcome knowledge and behavior was identified, possibly reflecting the ego-dystonic nature of OCD, with implications for the development of new behavioral and pharmacological interventions aimed at suppressing habits rather than focusing on dysfunctional beliefs.

## MATERIAL AND METHODS

### Participants

The study included 27 OCD patients and 27 controls, matched for relevant demographic variables (**Table S1**). Control subjects were recruited from the community; none of them was on psychiatric medication and they had never suffered from a psychiatric disorder. Patients were recruited through clinical referral from local psychiatric and psychological services or local advertisement. In addition, patients who participated in previous independent studies were contacted by phone. A consultant psychiatrist made DSM-5 diagnoses using an extended clinical interview, supplemented by the Mini International Neuropsychiatric Interview [42]. Patients were included if they met criteria for the diagnosis of OCD with no current comorbidity. Patients with OCD were not enrolled in the study if they scored less than 12 on the Yale–Brown Obsessive–Compulsive Scale (Y-BOCS) [43] and, in line with evidence that hoarding might represent a separate clinical entity [44], were excluded if they reported hoarding symptoms. Exclusion criteria for all participants were: current substance dependence, head injury, and current depression, indexed by Montgomery-Åsberg Depression Rating Scale exceeding 16 [45] during screening. Self-reported measures of anxiety were collected using the State-Trait Anxiety Inventory (STAI) [46]; and, in addition to Y-BOCS scores, self-reported measures of OCD symptomatology were collected using the Obsessive Compulsive Inventory-Revised (OCI-R) [47]. In patients, depression and anxiety symptoms were below the threshold for diagnosis of depressive or anxiety disorder (**Table S1**). 19 of the 27 patients were taking stable doses of serotonin reuptake inhibitor (SSRI) medication for a minimum of 8 weeks prior taking part in the study. As an adjunct to their SSRI, 3 of these patients were taking an antipsychotic (quetiapine). The remaining 8 patients were unmedicated, being either drug-naïve or off medication for at least 8 weeks prior taking part of the study. Most of the participants completed two other behavioural tasks, unrelated to the present study. The study was approved by the NHS East of England Cambridge Central Research Ethics Committee. Participants were reimbursed for their time and informed consent was obtained prior participation. No statistical methods were used to pre-determine sample size but our sample sizes are similar to those generally employed in the field, with power of 0.8 to detect effect sizes of 0.78 at α = 0.05, two-tailed.

### Procedure

Contingency degradation manipulation requires that the subject experiences the likelihood of the outcome given the presence or absence of a response. We adopted a free-operant, self-paced procedure whereby the participant could decide whether to press the space bar or not when presented with a white triangle on the screen. However, in a free-operant paradigm, the degree of contingency experienced can be determined partly by the behavior, and experienced contingency might in principle vary substantially across participants (e.g., someone who never responds would never experience P(O|A), and someone who never ceases responding would never experience P(O|∼A)). In schedules where reinforcer delivery is influenced by time (e.g. with a maximum reinforcer delivery rate or on an interval schedule), different subjects might experience similar reinforcer delivery rates despite different response rates. Therefore, we divided time into short 1 second interval (bin), and calculated ‘response’ versus ‘no response’ on a per-bin basis ensuring a close correspondence between programmed and experienced contingencies [16]. Accordingly, unbeknown to the participant each block was divided into bins, treated as a trial by the experimenter. The procedure was free-operant for the subject as trials were unsignaled and there was no inter-trial interval. In doing so, interpretation of our findings was not confounded by between-groups differences in experienced contingencies (Table 1).

### Experimental task

A white triangle permanently on the screen signaled the participant that he/she was free to press the space bar (or not press). When a reward was delivered, either following a key-press or not, a 25p image was shown at the end of the bin for 500 ms with the text “Reward, you win!” and a tone (**Figure 2A**). Upon each response, the triangle turned yellow until the end of the *a priori* specified bin to signal that a response has been recorded and prevent multiple responses within the same 1 second bin. If no outcome was delivered, no feedback was given and the next bin started. Note that if the participant did not respond for several time bins the white triangle stayed on the screen without anything else happening, unless a non-contingent reward occurred. A running total of pence accumulated within the block was displayed in the top right corner of the screen. There were 12 blocks, not explicitly labeled as such to the participants. However, at the beginning of each block the running total of pence was reset to 0, and at the end of each block causality judgments were collected on the relationship between pressing the key and receiving the 25p reward (**Figure 2A**). For each subject, the first 3 blocks (Blocks 1–3) were always presented in the same order (high contingency, degradation, extinction) providing an implicit training phase. The remaining 9 blocks (i.e., Block 4–12) were presented according to a Latin square design across participants (**Table 1**). Each block lasted for 2 minutes (120 unsignaled bins). If a response occurred during a given bin, the outcome was delivered at the end of the bin with probability P(O|A) defined *a priori* for that block; if no response occurred, the outcome was delivered with probability P(O|∼A) for that block (**Figure 2B**). Only the first space-bar press within the bin had any programmed consequences. The total number of responses within each bin was also recorded, but additional responding beyond the first response of the bin had no programmed consequences. The experiment was programmed using Psychtoolbox 3 [48–50]. The overall duration of the task was variable due to its free-operant nature, i.e. the rate of responding which was variable across participants determined the number of outcomes. In fact, we had a fixed amount of unsignalled bins for each block but delivery of a reinforcer delayed the start of the next bin. Hence the total duration depended also on the number of outcomes delivered but the average time for completion (34 minutes) did not differ between groups.

Our implementation of the task differed from previous ones available in the literature for some crucial aspects. Firstly, by using unsignalled time bins and by specifying the conditional probabilities *a priori* we ensured that experienced instrumental contingencies did not deviate substantially from the programmed ones. Secondly, in line with experimental studies in rodents where there is no explicit ‘punishment’ for responding we did *not* include a cost for responding (see Supplementary Material and Figure S5 for supporting results from pilot experiments with and without such costs). We found that introducing a cost induced a generalized reduction of responding, with no specific effect on determining responding in face of degradation (see Supplemental Material).

### Check on experienced contingency

In order to compute the experienced contingency for each subject for a given block, we recorded (i) the number of contingent outcomes (rewards delivered upon key press) (C1); (ii) the number of times that a key press was not associated with the delivery of an outcome (C2); (iii) the number of non-contingent outcomes (rewards delivered in the absence of a key press) (C3); (iv) the number of times that there was no key press and no outcome delivered (C4). We thus computed the experienced contingency based on the formula for contingency (ΔP) [16]:

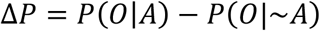

as:

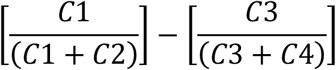

In very few instances experienced contingency could not be computed because there were no occurrences of either C1 and C2 or C3 and C4. In other words, the subject did not press the space bar throughout the block, or adopted a constant pressing rate with a consequential lack of no trials with no responses. However, in our entire data set (648 blocks; 12 blocks × 54 participants) this occurred only on 10 single occasions with 7 controls and 3 OCD patients adopting one of the specified strategies in one of the blocks during their experimental session. Inclusion or exclusion of these subjects did not affect the main findings, therefore, we retained data from these subjects for the analysis.

As expected, based on our implementation of the task, there was a very high correlation between the mean experienced contingency (based on experienced event frequencies) (Table 1) and the contingencies programmed a priori, for controls (r = 0.999, p < 0.001) and patients (r = 0.998, p < 0.001) alike. We therefore used the programmed contingencies for subsequent analysis. Importantly, the interpretation of our findings was not confounded by different levels of experienced contingencies between the two groups as no main effect of group (F_1,48.49_ = 0.01, p = 0.940) nor interaction between group and block (F_11,559.95_ = 1.06, p = 0.395) on experienced contingency was found.

As expected, there was a main effect of programmed contingency on the number of outcomes obtained (F_4,208_ = 38.831, p < 0.001). Even though OCD patients responded more at certain levels of instrumental contingencies, such increased behavior was not sufficient to lead to a higher number of obtained outcomes. In fact, there was no main effect of group on the number of outcomes obtained (F_1,52_ = 0.002, p = 0.960), nor a significant interaction between group and programmed contingency (F_4,208_ = 1.158, p = 0.330). These findings therefore rule out the possibilities that OCD patients’ behavior resulted in better outcomes overall or that OCD patients’ behavior was secondary to differences in reward rate. In addition, we used the BIS/BAS (Behavioral Inhibition System/Behavioral Approach System) questionnaire to measure reward responsiveness via the BAS reward responsiveness subscale [51]. Although data were available only for a subset of subjects (18 controls and 19 OCD) there was no group difference in reward responsiveness (t_35_ = 0.375, p = 0.710). There was no difference in response rate at the maximal contingency (Figure 3A, at ΔP = 0.6), but specifically for certain levels of contingency suggesting that the effect was due to reasons other than reward responsiveness.

### Data Processing and Analysis

All statistical tests were two-sided, and parametric or nonparametric testes applied as needed according to assumptions of the specific statistical test chosen. We analyzed performance in terms of response rate and causality judgements for different levels of instrumental contingency.

We adopted a two-step approach. Firstly, we identified if there was a difference between controls and patients in behavioral sensitivity to instrumental contingency. To this end we computed a response rate, obtained by dividing the number of responses by the number of bins for each block. For each dependent variable (response rate and causality judgement) programmed contingency was used as a within-subject factor and group as a between-subject factor (**Figures 3A, 3B** and **5A**). Data were collapsed across blocks having equal programmed contingencies. Analyses were performed in R version 3.3.1 (http://www.rproject.org/) using the ‘ez’ package for ANOVA. Levene’s test was used to verify homogeneity of variance. Mauchly’s test of sphericity was applied and Greenhouse–Geisser and Huynh–Feldt correction used for substantial (ε < 0.75) and minimal violation (ε≥0.75), respectively. To investigate the relationship between contingency judgments and response rate between groups, we used linear mixed-effects models (**Figure 3C**). Group was used as a fixed-effect factor; linear (and, where applicable, quadratic) causality judgments were used as continuous fixed-effect predictors. The maximal random effect structure justified by the design was specified [52] using mixed models [53].

Secondly, we tested specifically if behavior was habitual for those conditions in which we observed diminished sensitivity to instrumental contingency was observed in OCD patients and in which P(O|A) was stable and P(O|-A) was increased. Accordingly, we obtained a ratio score by considering pairs of contingent and corresponding degraded blocks [17]: for each pair, the number of responses in the contingent block was divided by the sum of responses in both the contingent and degraded blocks. Thus the ratio score represents the number of responses in the contingent condition as a proportion of the total responses made across both contingent and degraded condition, with values close to 1 indicating high sensitivity to contingency and values close to 0.5 indicating habitual behavior. Homogeneity of variance across groups was verified via Levene’s test and Student’s t-test applied accordingly (**Figures 4A-4C**). Data collection and analysis were not performed blind to the conditions of the experiment.

## ACKNOWLEDGMENTS

We thank all the patients and volunteers who took part in this study. We thank Anthony Dickinson and Simon White for their advice. This research was funded by a Wellcome Trust Senior Investigator Award (104631/Z/14/Z) to T.W. Robbins. Work was completed at the Behavioural and Clinical Neuroscience Institute, University of Cambridge, Cambridge, UK, supported by a joint award from the Medical Research Council and Wellcome Trust (G00001354). M.M. Vaghi is supported by a Pinsent Darwin Scholarship in Mental Pathology and Angharad Dodds John Bursary in Mental Health and Neuropsychiatry. The work was previously presented in part at Society for Neuroscience, San Diego, CA, 2016.

